# Quantitative proteomics uncovers a role for *REVEILLE8-*like clock genes in osmotic and salt stress response in Arabidopsis

**DOI:** 10.64898/2026.06.08.730740

**Authors:** LE Grubb, I Khodabocus, S Scandola, D Mehta, RG Uhrig

## Abstract

The plant circadian clock governs the precise regulation of plant developmental and environmental responses within a 24 h photoperiod. Consisting of a series of interconnected transcription factor moderated feedback loops, each operating at specific times of day within the photoperiod, more than 30% of *Arabidopsis* genes show circadian regulation. *REVEILLE* (RVE) genes exhibit highest expression in the afternoon, and act as activators of afternoon-expressed clock genes, including *TOC1* and *PRR5*. Specifically, RVE4, 6 and 8 have been identified as regulators of both plant growth and development, along with temperature responses. Previous work using Arabidopsis has implicated soybean RVE8-like proteins in drought responses, however, there remains a lack of understanding of how these RVEs control abiotic stress responses, particularly relating to proteome-level regulation. Here, using quantitative proteomics, we resolve how the *rve4 6* 8 proteome responds to osmotic and salt stress at end-of-day and end-of-night timepoints. Our results indicate that *rve4 6 8* plants have substantially altered regulation of proteins related to osmotic stress, photosynthesis and phenylpropanoid metabolism, particularly at end-of-day. Further, we resolve new drought-responsive targets impacted by the loss of RVE4, 6, and 8 that are involved in vesicle transport, fatty acid metabolism and abscission. Overall, our data provide a new resource for understanding how RVE8-like proteins impact plant drought-like responses, offering opportunities for future chronoculture-informed breeding of climate-resilient crops.

## INTRODUCTION

For global agriculture, the severity and prevalence of abiotic stressors, such as drought and salinity, are increasing drastically with the changing climate (Cogato et al., 2019). Drought and salinity induce cellular osmotic stress, which can reduce plant water uptake, ultimately leading to ion imbalance, oxidative damage, and cell death (Choudhury et al., 2017). In addition, plants typically close their stomata upon perception of these stressors, resulting in disrupted photosynthesis (Yang et al., 2021), and serious impacts on plant growth, development, and overall yield.

The plant circadian clock is critical for the coordination of appropriate daily (diel) plant responses, regulating growth and development, as well as responses to a variety of internal and external stimuli (Greenham and McClung, 2015; Nohales and Kay, 2016; McClung, 2019). In *Arabidopsis thaliana* (Arabidopsis), at least 30% of plant genes have been shown to be under direct circadian control (Covington et al., 2008), many of which are involved in plant responses to abiotic stressors (Simon et al., 2020; Kidokoro et al., 2021). As plants experience peak exposure to drought stress during the day, rhythmic expression of drought-inducible factors may allow for maximal response at the optimal time in the diurnal cycle (Grundy et al., 2015). In addition, genes involved in stress-responsive plant hormone metabolism, such as ABA, have been shown to have rhythmic expression (Seung et al., 2012; Adams et al., 2018), with the expression of ABA biosynthesis and signaling response genes having been shown to be under the control of the circadian clock genes *TIMING OF CAB EXPRESSION 1* (*TOC1*) (Legnaioli et al., 2009) and *LATE ELONGATED HYPOCOTYL* (*LHY)* (Adams et al., 2018). Several other Arabidopsis circadian clock genes have also been implicated in the plant response to osmotic stress including *GIGANTEA* (*GI)* (Liu et al., 2024), the evening complex genes, *EARLY FLOWERING 3* (*ELF3*), *ELF4,* and *LUX ARRHYTHMO (LUX)* (Tang et al., 2025), and *PSEUDO-RESPONSE REGULATORS 5* (*PRR5*), *PRR7* and *PRR9* (Prasetyaningrum et al., 2023). Therefore, understanding how the circadian clock intersects with plant abiotic stress responses could allow for chronoculture-informed approaches to the generation of climate-resilient crops (Steed et al., 2021). While many of these studies identify a role for the circadian clock using transcriptome data, it is well known that transcriptome changes are often disconnected from functional temporal and spatial proteome responses (Hassan et al., 2025; Mehta et al., 2025; Hassan et al., 2026), rendering it important to understand circadian clock mediated processes at the protein level.

The REVEILLE (RVE) 8-like MYB transcription factors, consisting of RVE8 and its homologs, RVE4 and RVE6, have been identified as activators of the core plant circadian clock (Farinas and Mas, 2011; Rawat et al., 2011; Hsu et al., 2013). RVE8 directly binds the promoter region of *TOC1* and *PRR5* (Farinas and Mas, 2011; Rawat et al., 2011), and has also been shown to induce expression of the evening complex, consisting of *ELF3, ELF4,* and *LUX* (Hsu et al., 2013). Both the *rve8* (Farinas and Mas, 2011; Rawat et al., 2011) and *rve4 6 8* mutants possess an elongated circadian period (Hsu et al., 2013), with *rve4 6 8* plants exhibiting larger overall plant size, resulting from a faster growth rate, larger average cell size, and delayed flowering time (Gray et al., 2017). In addition to roles in growth and development, RVE8 is important for the diel accumulation of anthocyanin, promoting expression of anthocyanin biosynthesis genes around dawn in Arabidopsis (Pérez-García et al., 2015), pear (Li et al., 2020; Li et al., 2024), and crocus (Bhat et al., 2023). As well, roles for RVE8-like proteins in abiotic stress responses have been emerging. Specifically, RVE4 and RVE8 have been identified to regulate early heat-shock induced gene expression (Li et al., 2019), and function in the plant cold stress response, through activation of *DEHYDRATION RESPONSIVE ELEMENT (DRE) BINDING PROTEIN 1* (*DREB1*) (Kidokoro et al., 2021; Kidokoro et al., 2023). Thus RVE8-like proteins appear to be key transcriptional regulators of plant temperature responses.

Using a multi-omics approach, we recently reported time-of-day differences in several important cell processes for the *rve4 6 8* mutant (Scandola et al., 2022). These include altered carbohydrate metabolism leading to starch excess at dawn, as well as reduced proteasome activity and altered amino acid, organic acid, and lipid metabolism, which may explain observed phenotypic growth differences relative to the wild-type Col-0 (Col-0) (Scandola et al., 2022). In addition, metabolic differences indicated a potential role for the RVE8-like proteins in plant responses to osmotic stress (Scandola et al., 2022). Key metabolome differences in *rve4 6 8* plants included increased levels of mannose, which functions in stomatal conductance (Li et al., 2016), and alpha-linolenic acid, which plays a role in cell membrane fluidity (Upchurch, 2008). Notable proteome changes in *rve4 6 8* plants included a reduction in jasmonic acid (JA) signaling protein abundance, which is linked to alpha-linolenic acid (Ruan et al., 2019), alongside an increased abundance of sulfur assimilation proteins, which are critical for production of abscisic acid (ABA) and glutathione; both important metabolites for plant drought and salt stress responses (Kopriva, 2006). In addition to identifying likely roles for RVE8-like proteins in drought and salinity, a recent study found Arabidopsis expressing soybean RVE8 to have increased tolerance to drought and salt stress (Bao et al., 2024), further implicating the RVE8-like proteins as important for these osmotic stress responses.

With these findings suggesting RVE8-like proteins play an important role in osmotic stress responses, there remains a lack of molecular information resolving the biological process underpinning this role. Here, using a quantitative proteomic approach, we address this with time-of-day precision by resolving the proteome differences in *rve4 6 8* seedlings exposed to osmotic and salt stress conditions. Our results implicate RVE8-like proteins in the regulation of a variety of stress responsive biological processes, including anthocyanin, photosynthesis, water deprivation response and ABA signaling, among others. We also provide validation for the involvement of RVE8-like proteins in these processes through the physiological analysis of plant genetics corresponding to proteins exhibiting a significant change in their protein abundance. Given the association of RVE8-like proteins with multiple agronomically important traits, such as plant biomass, carbohydrate metabolism and thermotolerance, it is important to understand their function under drought-like conditions for potential application in developing chronoculture-informed climate resilient crops.

## RESULTS

### The rve4 6 8 mutant has decreased tolerance for osmotic and salinity stress

Given the increased tolerance conferred by soybean RVE8 when expressed in Arabidopsis (Bao et al., 2024), along with our previous proteome and metabolome results suggesting that Arabidopsis RVE8-like proteins have a role in drought-like stress responses (Scandola et al., 2022), we sought to further elucidate the role of Arabidopsis RVE8-like proteins in osmotic and salt stress responses. We first examined the germination and primary root length of *rve4 6 8* seedlings upon exposure to osmotic stress conditions over 10 days. For osmotic stress responses, we germinated seeds on 0.5x MS plates containing 300 mM mannitol or transplanted seedlings to observe primary root length at 8 days post-transplant on 0.5x MS containing 50 mM, 100 mM, or 200 mM mannitol. We observed a significantly lower germination rate for *rve4 6 8* seedlings (**Figure 1A, B, D**) and a significantly shorter primary root under the 50 mM mannitol condition (**Figure 1F, H**), suggesting that RVE8-like proteins play a substantive role in Arabidopsis response to osmotic stress. Next, we examined germination and primary root growth of *rve4 6 8* seedlings under salt stress conditions, using 0.5x MS containing 100 mM NaCl for germination assays and 25 mM, 50 mM, and 100 mM NaCl for root growth experiments. Similar to the mannitol treatment, *rve4 6 8* seedlings showed a significantly lower germination rate (**Figure 1A, C, E**) and shorter primary roots (**Figure 1G, H**) under salt stress. Overall, our drought-like stress phenotyping results indicate an important role for RVE8-like proteins in response to osmotic and salinity stress.

**Figure 1.**
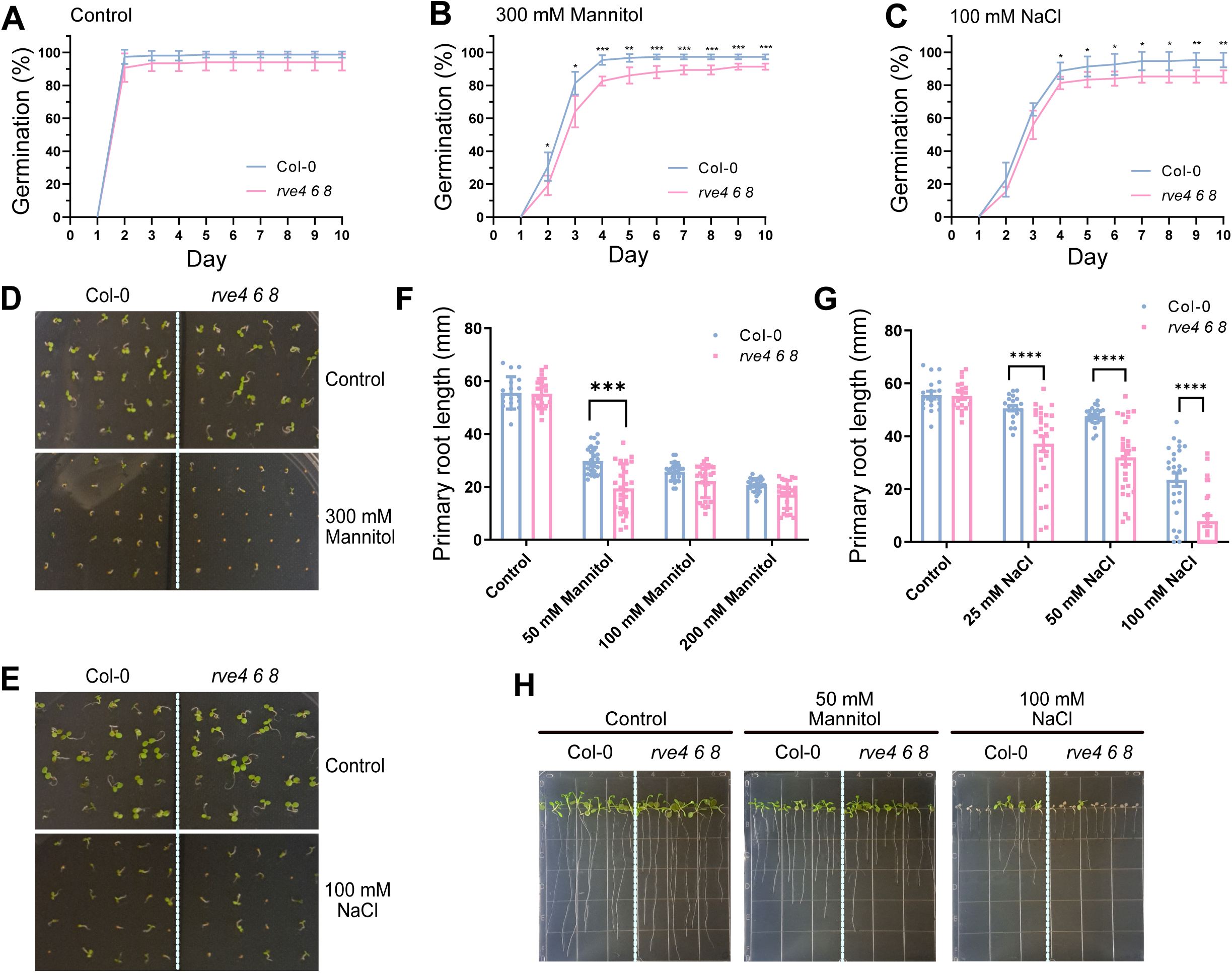
The *rve4 6 8* mutant is susceptible to drought and salt stress. **A-C**) Germination rate (%) of WT and *rve4 6 8* grown for 10 days on 0.5x MS medium control (**A**), supplemented with 300 mM Mannitol (**B**) or 100 mM NaCl (**C**). Data show mean and SD of 5 independent plates. Asterisks indicate significant differences (Unpaired t-test; *p<0.05; **p<0.01; ***p<0.001). **D**) Germination phenotype of WT and *rve4 6 8* on 0.5x MS plates at 4 days after plating under Control or 300 mM Mannitol conditions. **E**) Germination phenotype of WT and *rve4 6 8* on 0.5x MS plates at 4 days after plating under Control or 100 mM NaCl conditions. **F-H**) Primary root length analysis of WT and *rve4 6 8* transferred to 0.5x MS medium supplemented with 0, 50, 100 or 200 mM Mannitol (**F, H**) or 0, 25, 50 or 100 mM NaCl (**G, H**) analyzed at 8 days following transfer to experimental condition. Data are mean with SEM (n= >18). Asterisks indicate significant differences (Unpaired t-test; ***p<0.001; ****p<0.0001). **H**) Photographs taken 8 days following transfer to 50 mM Mannitol and 100 mM NaCl.

### Quantitative proteomic analysis identifies proteome changes under osmotic and salt stress in rve4 6 8

To further elucidate the molecular role of RVE8-like proteins under osmotic and salt stress conditions, we performed a quantitative proteomics analysis to determine changes in the Arabidopsis seedling proteome. As RVEs are key activators of the circadian clock, we sampled at two different timepoints for a time-of-day perspective: Zeitgeber time 0 (ZT0), also referred to as end-of-night (EN), and ZT12, referred to as end-of-day (ED). Based on our growth experiments, we chose to germinate seeds on 0.5x MS control media and transferred seedlings to mannitol (50 mM) or NaCl (100 mM) 5 days post-imbibition, followed by harvesting whole seedlings after 8 days on stress media (**Figure 2A**). Across our experiments, we quantified 6,630 protein groups (**Supplemental Table 1**). Using this data, we next determined proteins that were differentially abundant in *rve4 6 8* vs. Col-0 under our control, osmotic and salt conditions at ZT0 (**Figure 2B**) and ZT12 (**Figure 2C**; Log_2_FC > 0.58 or < -0.58; *q-value* < 0.05). Similar to our previous time-course analysis of Arabidopsis shoots and roots under osmotic and salt stress conditions (Rodriguez Gallo et al., 2023), we noted that there were only 2 overlapping differentially abundant proteins between osmotic and salt treatments at ZT0 (**Figure 2B**) and only 8 overlapping proteins between osmotic and salt treatment at ZT12 (**Figure 2C**). Overall, we found the control condition to have the greatest number of differentially abundant proteins and salt treatment the least at ZT0 (**Figure 2B**), with the inverse being true for ZT12 (**Figure 2C**). These results underscore the importance of time of day when undertaking stress experiments.

**Figure 2.**
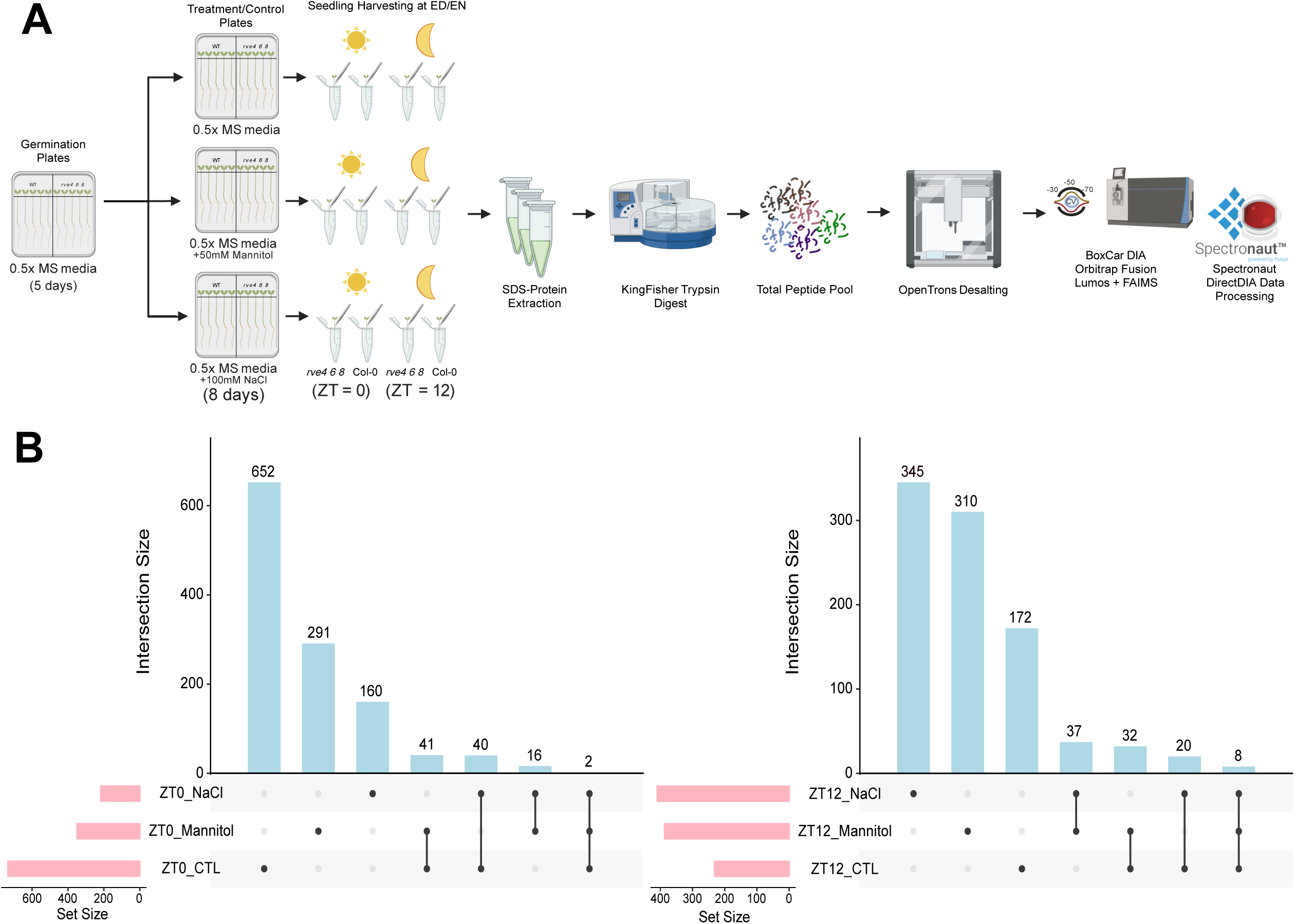
Analysis of proteome changes in *rve4 6 8* under mannitol and salt stress. **A**) Depiction of the quantitative proteomic workflow. **B-C**) Upset plots of significantly changing proteins (Log2FC >0.58 or <-0.58) in *rve4 6 8* at ZT0 (**B**) and ZT12 (**C**).

To delve further into which biological processes are differentially regulated in *rve4 6 8* under osmotic and salt stress conditions, we performed a gene ontology (GO) enrichment analysis. As there are inherent proteome differences between *rve4 6 8* and Col-0 plants, we subtracted the Log_2_FC of DAPs from the control condition from the Log_2_FC of DAPs under treatment conditions and used a differential threshold of Log_2_FC < -0.58 or > 0.58 for further analyses (**Figure 3**). Interestingly, we observe more enriched GO terms under both osmotic and salt stress treatments at ZT12, suggesting greater divergence in stress-responses of *rve4 6 8* and Col-0 at end-of-day versus end-of-night. Given the peak midday abundance of RVE8 protein (Rawat et al., 2011), this was not completely surprising. Here, we find the *rve4 6 8* stress proteome to be enriched in biological processes related to drought-like stress responses, including ‘*Response to Water Deprivation*’ (GO:0009414) and ‘*Response to Water*’ (GO:0009415) under both osmotic and salt stress conditions (**Figure 3**). In response to abiotic stress, such as osmotic or salt stress, plants produce reactive oxygen species (ROS) (Ali et al., 2023). In line with this, we observe enrichment of GO terms ‘*Toxin Metabolic Process*’ (GO:0009404), ‘*Reactive Oxygen Species Metabolic Process*’ (GO:0072593), and ‘*Detoxification*’ (GO:0098754), particularly at ZT12 under the osmotic stress condition (**Figure 3**). Interestingly, we observed the highest enrichment of the GO terms ‘*Photosynthesis, Light Reaction*’ (GO:0019684) and ‘*Electron Transport Chain*’ (GO:0022900) under the salt treatment at ZT12, suggesting potentially altered regulation of photosynthetic processes in *rve4 6 8* plants under salt stress.

**Figure 3.**
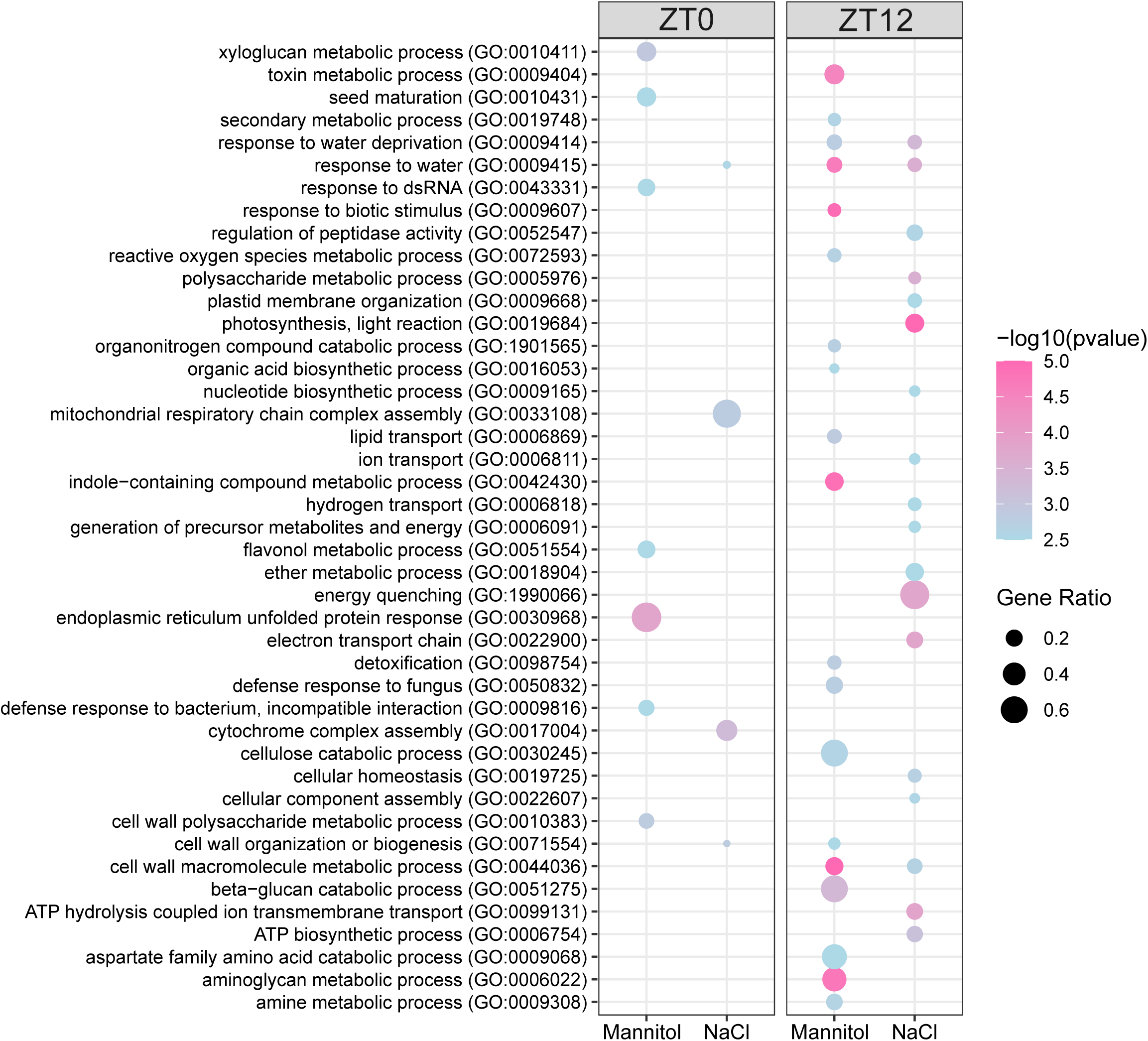
Gene Ontology enrichment analysis. Dotpot representation of enriched biological processes using significantly changing proteins from *rve4 6 8*. Differentially abundant proteins (DAPs) in *rve4 6 8* at ZT0 and ZT12 under control conditions were subtracted from Mannitol and NaCl stress DAPs to account for genetic background differences between Col-0 and *rve4 6 8*. Only DAPS with a control-condition normalized Log2FC > 0.58 or < -0.58 following this subtraction were considered. Node size indicates gene ratio (number of proteins in the condition / total quantified proteins in the study). Blue to pink colouration indicates the -log10 (p-value) for the indicated biological process. Only GO categories with p-value < 0.01 and total family size < 500 are included.

To further contextualize the proteome differences between *rve4 6 8* and Col-0 under osmotic and salt stress conditions, and to resolve whether these biological processes might be up- or down-regulated, we performed an association network analysis using stress-specific, significantly changing DAPs (**Figure 4**). Similar to our GO analysis, we also observe clusters of proteins related to water deprivation response in both the osmotic and salt treatment. Interestingly, many of these proteins appear to be of higher abundance in *rve4 6 8*, suggesting disrupted regulation of these stress-responsive proteins at our sampling time. We additionally observed higher abundance of other stress-related biological processes in *rve4 6 8* including glutathione metabolism, phenylpropanoid metabolism, detoxification, and vesicle trafficking. Strikingly, under the salt treatment, there were a high abundance of DAPs related to starch and sucrose metabolism, which may be related to the starch excess phenotype previously reported (Scandola et al., 2022), lipid metabolism, photosynthesis, and chlorophyll metabolism, particularly at ZT12 (**Figure 4B**).

**Figure 4.**
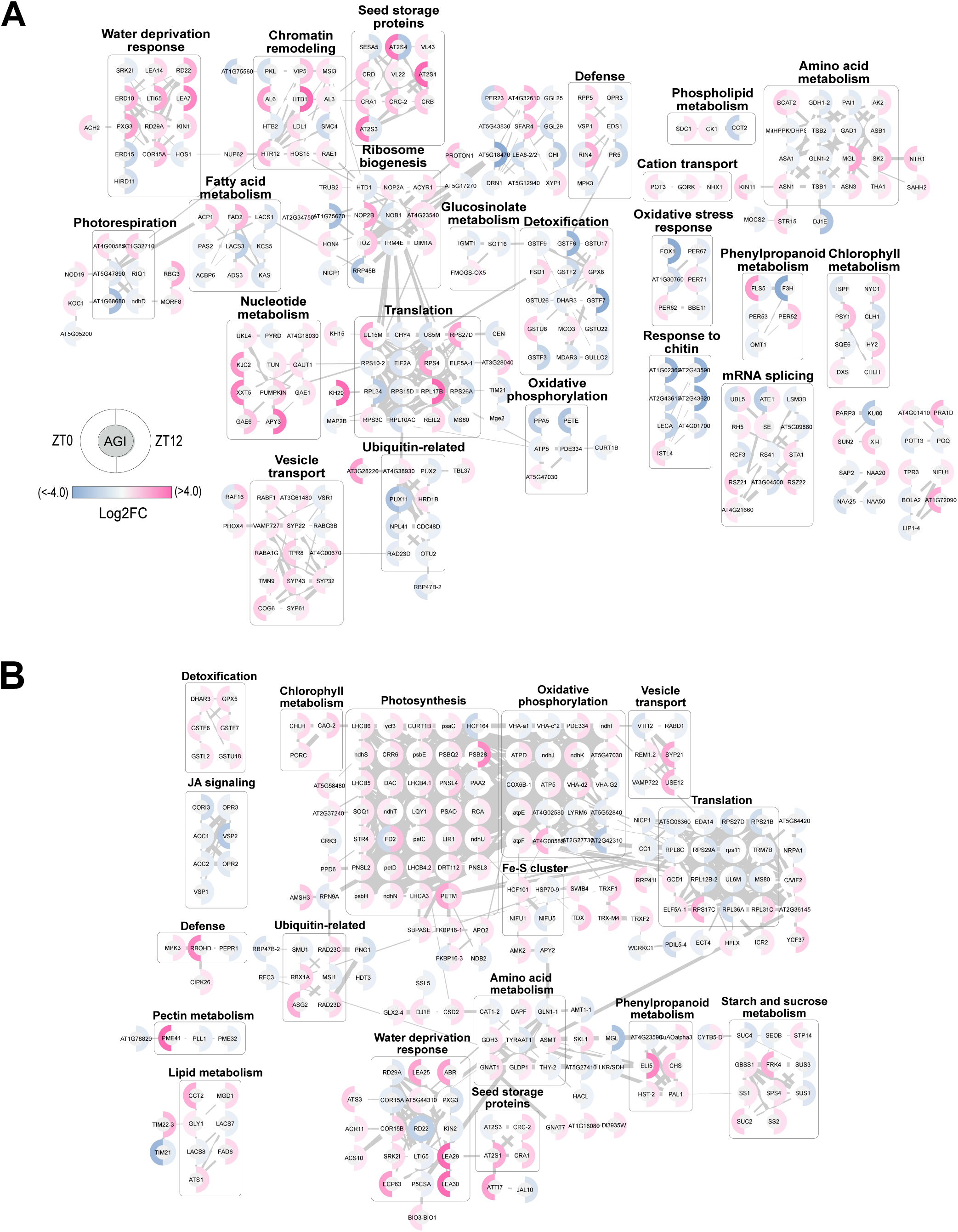
Association network analysis of proteome differences in *rve4 6 8* under mannitol (A) and salt stress (B). Association network depicting significantly changing protein abundance between *rve4 6 8* under mannitol (**A**) and salt stress (**B**). Differentially abundant proteins (DAPs) in *rve4 6 8* at ZT0 and ZT12 under control conditions were subtracted from Mannitol and NaCl stress DAPs to account for genetic background differences between Col-0 and *rve4 6 8*. Only DAPS with a control-condition normalized Log2FC > 0.58 or < -0.58 following this subtraction were considered. Networks were produced using Cytoscape with the STRING-DB database and enhancedGraphics app (see Materials and Methods) using all datatypes and an edge score threshold of > 0.7. Nodes with no edges or edges < 0.7 were removed and only nodes connected in a group of more than 2 are shown. Grey boxes indicate biological process relationships among the nodes. This was determined using the Cytoscape Functional Annotation tool.

To determine metabolic pathways impacted by RVE8-like proteins under osmotic or salt stress conditions, we next used the Plant Metabolic Network (PMN; https://pmn.plantcyc.org) to examine enriched of metabolic processes among the DAPs in *rve4 6 8* relative to Col-0, using only DAPs unique to the stress treatments (**Supplemental Table 2**). Here we found several enriched pathways that corroborate both our association network analysis and GO enrichment. This includes metabolic pathways associated with fatty acid biosynthesis, jasmonic acid biosynthesis, photosynthesis, and starch biosynthesis (**Supplemental Table 2**). Overall, these results implicate RVE8-like proteins in a variety of stress-responsive and developmental processes in response to osmotic and salinity stress.

We next investigated which proteins were significantly different between ZT12 and ZT0 in both Col-0 and *rve4 6 8* under control, osmotic, or salt treatments to understand whether the observed proteome changes in the mutant are due to altered time of day abundance (**Supplemental Figure 1**). Under all conditions, we find lower overall DAPs in *rve4 6 8* and very few proteins differentially regulated at end-of-day vs. end-of-night in both genotypes. Under control conditions, there were 17/239 (∼7%) proteins differentially regulated by timepoint in *rve4 6 8*, with 40/377 (∼11%) under mannitol treatment and 38/269 (14%) in salt treatment. We then performed GO enrichment on the proteins that were differentially regulated at ZT12 vs. ZT0 in Col-0, but not in *rve4 6 8* as there were too few DAPs. Under control conditions, we observed an enrichment of ‘*Abscisic Acid-Activated Signaling Pathway*’ (GO:0009738), ‘*Chromatin Organization*’ (GO:0006325), ‘*Proteolysis*’ (GO:0006508), ‘*Vesicle-mediated Transport*’ (GO:0016192), and ‘*Brassinosteroid Mediated Signaling*’ (GO:0009742). Under osmotic stress conditions, we find ‘*Detoxification*’ (GO:0098754), ‘*Response to Salt Stress*’ (GO:0009651), ‘*Response to Oxidative Stress*’ (GO:0009679), ‘*Circadian Rhythm*’ (GO:0007623), ‘*Lignin Biosynthetic Process*’ (GO:0009809), ‘*Response to Abscisic Acid*’ (GO:0009737), ‘*Response to Cold*’ (GO:0009409) and ‘*Response to Mannitol*’ (GO:0010555), while under salt stress we observe an enrichment of biological processes related to ‘*Cell Wall Organization*’ (GO:0071555), ‘*Photosynthesis*’ (GO:0015979), ‘*Phenylpropanoid Biosynthetic Process*’ (GO:0009699), ‘*Response to Abscisic Acid*’ (GO:0009737), ‘*Vesicle-mediated Transport*’ (GO:0016192), ‘*Response to Cold*’ (GO:0009409), ‘*Response to Oxidative Stress*’ (GO:0009679), and ‘*Starch Biosynthetic Process*’ (GO:0019252). Thus, it appears that many of the processes we attribute to *rve4 6 8* under drought-like conditions may be due to altered timing of protein expression or a loss of peak abundance.

### Loss of RVE4 6 8 alters photosynthesis, carbohydrate metabolism and anthocyanin production in response to salt stress

Our GO analysis identified an enrichment of photosynthetic processes among *rve4 6 8* DAPs under salt stress (**Figure 3**), indicating a potential role for RVE8-like proteins in modulating photosynthesis upon exposure to salt. This was further resolved by our association network to indicate increased abundance of these photosynthetic proteins in *rve4 6 8* at ZT12 relative to Col-0 (**Figure 4B**). To validate these proteomic observations, we grew Col-0 and *rve4 6 8* plants on soil and after 21 days, followed by the application of water (control), or 300 mM NaCl for 7 days (**Figure 5A**). We measured a suite of photosynthetic parameters using MultispeQ (Kuhlgert et al., 2016) at ZT3, the time of peak RVE8 protein abundance (**Figure 5B-C; Supplemental Figure 2**) (Rawat et al., 2011). Here, we find Phi2, or quantum efficiency for photosystem II photochemistry (Kuhlgert et al., 2016), to be significantly increased in Col-0 upon salt treatment, however it is already high and unchanging upon salt treatment in *rve4 6 8* (**Figure 5B**). Phi2 represents the amount of light received by the leaf that goes to photosystem II, which is ultimately used to drive CO_2_ fixation into carbohydrates (Kuhlgert et al., 2016). The significantly higher Phi2 observed in *rve4 6 8* plants under control conditions may relate to the previously reported starch excess phenotype (Scandola et al., 2022). Another photosynthetic parameter of note was PhiNO, which is the basal dissipation of light energy (Kuhlgert et al., 2016). In salt-treated treated *rve4 6 8* plants, we found a higher PhiNO than similarly treated Col-0 (**Figure 5C**). This represents the amount of light that has not been used for photosynthesis or dissipated, which can result in photosystem damage (Kuhlgert et al., 2016). This suggests higher light-induced stress in *rve4 6 8* plants relative to Col-0 under salt stress.

**Figure 5.**
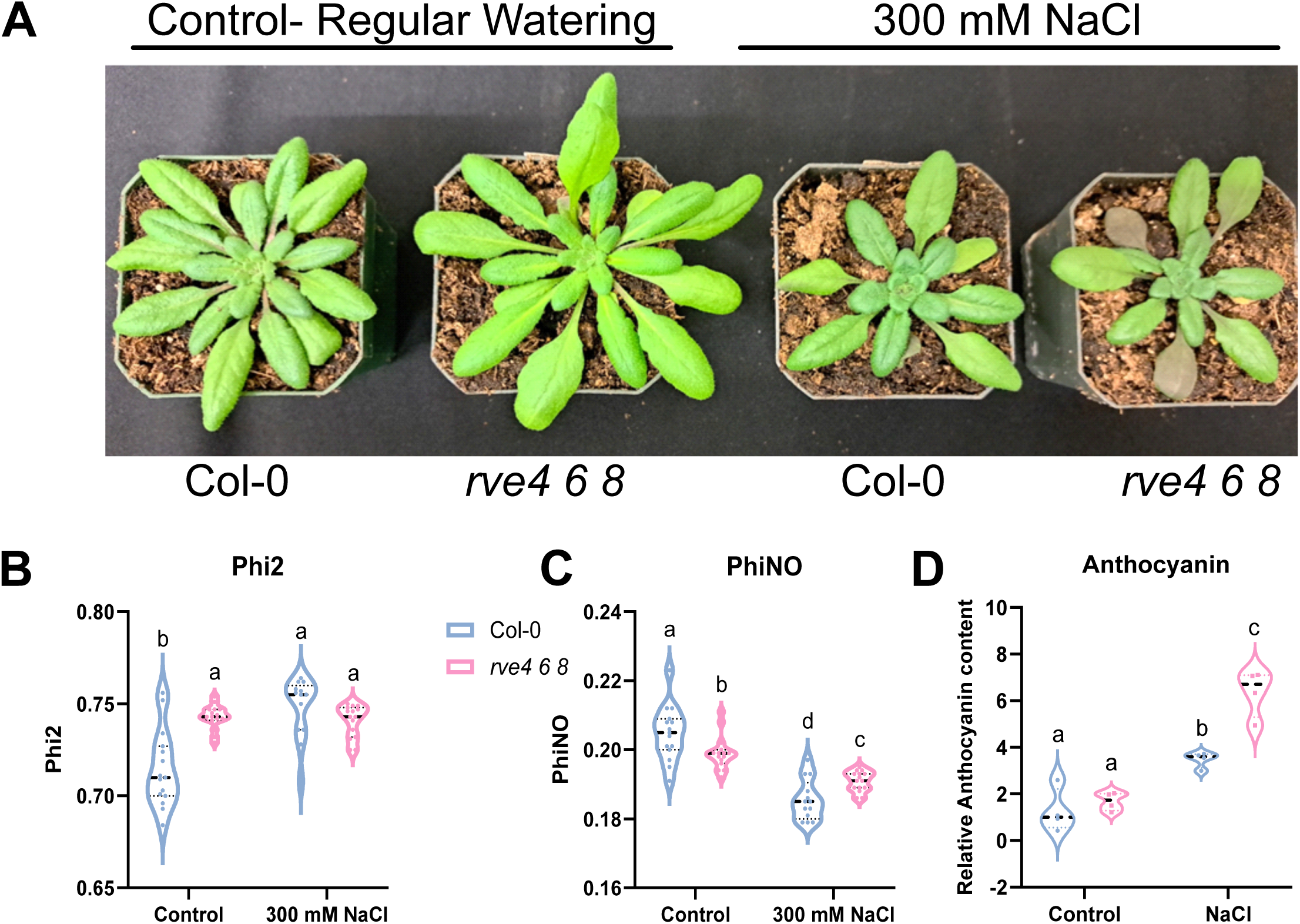
*rve4 6 8* shows differences in photosynthetic parameters and anthocyanin production under salt stress. **A**) Representative images of Col-0 and *rve4 6 8* mutant plants grown under control conditions with regular watering (Control) or watering with 300 mM NaCl (see Materials and Methods). All plants were grown for 21 days with regular watering prior to treatment for 7 days. Pictures were taken 7 days following the beginning of treatment. **B-C**) Violin plots indicating photosynthetic parameters of the plants described in (A) showing Phi2 (**B**) and PhiNO (**C**) for Col-0 (blue) and *rve4 6 8* mutant (pink) under control and 300 mM NaCl. **D**) Violin plots indicating anthocyanin content of the plants described in (**A**). Violin plots indicate the median and n=16 for photosynthesis measurements and 4 for anthocyanin. Letters denote statistical significance (p < 0.05) based on a 2-way ANOVA followed by Fisher’s LSD test for multiple comparisons.

The increased stress level of *rve4 6 8* was also phenotypically evident in the Arabidopsis plants (**Figure 5A**), where, as under control conditions, *rve4 6 8* plants were larger than Col-0 (Gray et al., 2017). However, this was not the case for salt-watered plants, where *rve4 6 8* plants appeared more purple. This leaf purpling was attributed to significantly greater amounts of anthocyanin in *rve4 6 8* relative to Col-0 (**Figure 5D**). As anthocyanins play a key role in plant response to photooxidative stress (Li and Ahammed, 2023), this indicates that *rve4 6 8* plants are experiencing greater photooxidative stress under salt stress conditions relative to Col-0. Thus, the results suggest that photosynthesis and general stress responses, including anthocyanin production, are dysregulated in *rve4 6 8* plants in response to salinity stress, consistent with our underlying proteome findings.

### Validation of putative targets in response to osmotic and salt stress

To determine whether our proteomic analysis identified some proteins that may be regulated by RVE8-like transcription factors in response to osmotic and salt stress, we selected several targets for phenotypic validation that were differentially abundant in *rve4 6 8* relative to Col-0 in our proteomic analysis under one or both stress conditions. Additionally, we triaged the DAPs for candidates that maintain rhythmic gene expression patterns, which was done using the Diurnal DB (https://diurnal.mocklerlab.org/) database. We ordered homozygous T-DNA insertion knockouts from the Arabidopsis Biological Resource Centre (ABRC) and germinated them on 0.5x MS media prior to transplanting to 0.5x MS plates containing either 50 mM mannitol (osmotic stress) or 100 mM salt (salinity stress) after 5 days of growth (8 days post-imbibition (**Figure 6**). Primary root length was then assessed after 8 days on stress treatment plates. Of the gene-deficient lines tested, we observed significantly different root lengths on our salt stress condition, but not our chosen mannitol condition.

**Figure 6.**
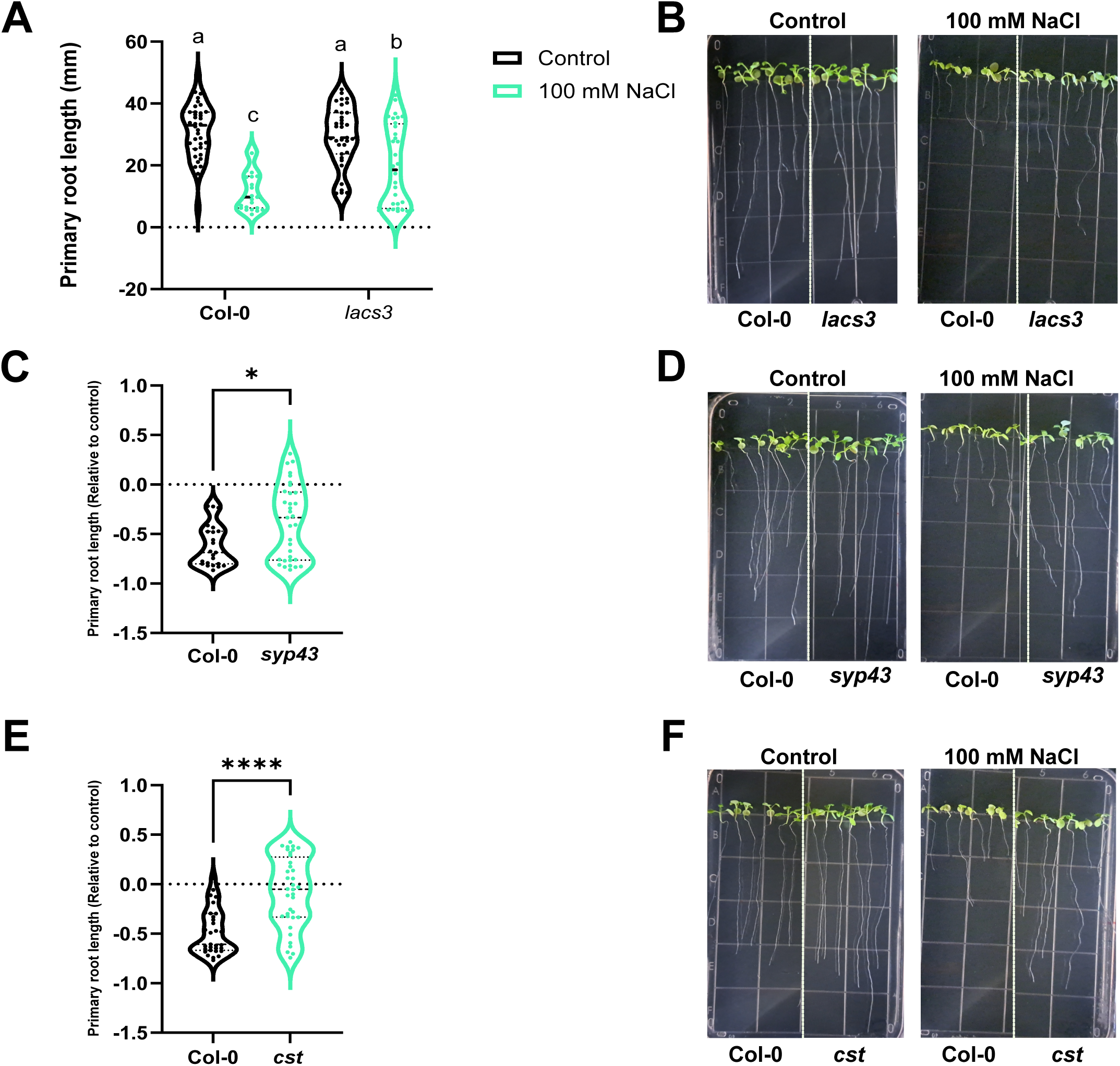
Root growth phenotypic validation of candidate lines under salt stress. Root lengths were measured for Col-0 and *lacs3* (**A-B**), *syp43* (**C-D**) or *cst* (**E-F**) loss-of-function seedlings following 8 days treatment on Control or 100 mM NaCl plates. A 2-way ANOVA followed by Fisher’s LSD test for multiple comparisons was performed for *lacs3*(**A**). For *syp43* (**C**) and *cst* (**E**) data were normalized to control conditions followed by use of a non-parametric Mann-Whitney test. Letters denote statistical significance (p<0.05; **A**), asterisks indicate statistical significance (**C, E**), *, p < 0.05; ****, p < 0.0001, n = at least 24 plants per genotype. Line in middle of violin plot indicates median. *LONG-CHAIN ACYL-COA SYNTHETASE3* (*LACS3*; AT1G64400), *SYNTAXIN OF PLANTS 43 (SYP43*; AT3G05710), and *CAST AWAY* (*CST;* AT4G35600).

**Figure 7.**
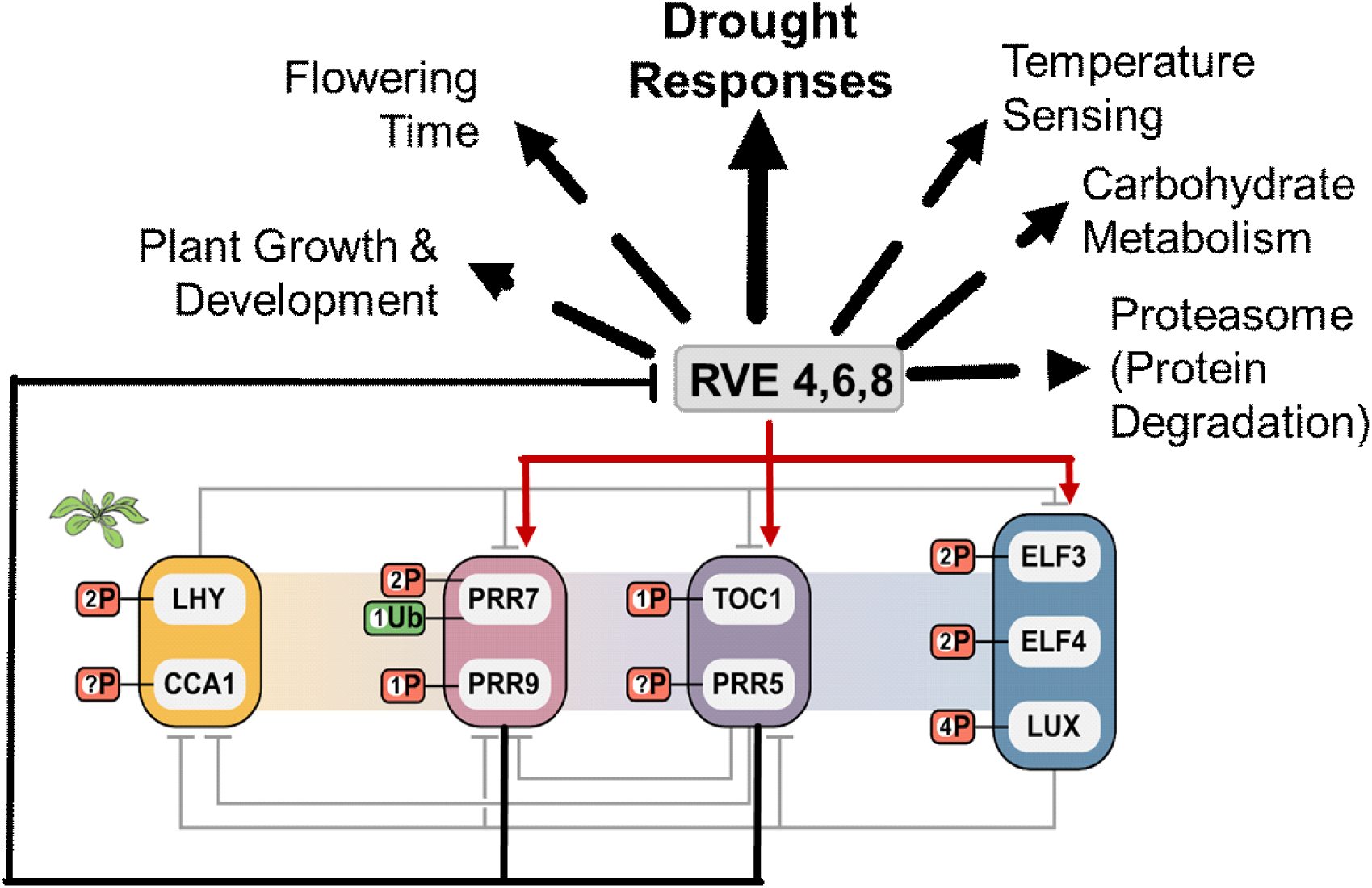
Summary of the roles of RVE8-like proteins in growth, development, and abiotic stress responses

With our association network analysis identifying fatty acid biosynthesis as a commonly differentially regulated process in both our osmotic and salt treatment, we selected LONG-CHAIN ACYL-CoA SYNTHETASE 3 (LACS3; AT1G64400) for further analysis (**Supplemental Table 1**). Subsequent querying of the Bio-analytic Resource (BAR; ePlant, https://bar.utoronto.ca/eplant/) supported this choice, as an induction of *LACS3* expression under osmotic salt stress was found. Further, *LACS3* was also recently found to be differentially expressed under drought stress (Ghanbari Moheb Seraj et al., 2025) and salt stress (Xu et al., 2024), but with no prior relationship to the circadian clock. Here, our phenotypic root growth analyses of *lacs3* plants found a greater tolerance to 100 mM salt relative to Col-0, maintaining a longer primary root length under salt stress in comparison to Col-0 (**Figure 6A-B**). Plant vesicle trafficking is also known to be responsive to abiotic stressors (Moura et al., 2025), with our proteomics results finding a greater abundance of SYNTAXIN OF PLANTS 43 (SYP43; AT3G05710) in *rve4 6 8* plants under osmotic stress (**Supplemental Table 1**). This syntaxin has not previously been characterized under any drought-like stress condition, however the BAR database indicated increased expression of *SYP43* under both osmotic and salt stress. Here, we find *syp43* plants to exhibit an increased tolerance to salt stress relative to Col-0 (**Figure 6C-D**), indicating that SYP43 may be a potential drought-stress susceptibility factor. Finally, we selected a receptor-like cytoplasmic kinase (RLCK), CAST AWAY (CST; AT4G35600), which has a previously indicated role in abscission (Burr et al., 2011; Groner et al., 2016). The process of abscission is intricately linked to ABA signaling, which is important for osmotic stress response (Gómez-Cadenas et al., 2002). CST was one of the DAPs with the largest change in abundance we observed under both osmotic and salt treatments (**Supplemental Table 1**), with greater abundance in *rve4 6 8* at ZT12 under osmotic stress, and ZT0 under salt stress. Our phenotyping assays of *cst* plants revealed a decreased susceptibility to salt stress relative to Col-0, but no difference under mannitol (**Figure 6E-F**). Our lack of osmotic stress-induced changes may indicate need for a higher mannitol concentration. Our choice of 50 mM mannitol was based on our phenotyping results for *rve4 6 8,* where this was the only concentration in which we observed a significant difference in primary root length. Taken together, our phenotypic results provide experimental validation for several candidates identified in our proteomic analysis and further connect fatty acid metabolism, vesicle trafficking and ABA-related biological processes (e.g., abscission) with RVE8-regulated plant responses to abiotic stress.

## DISCUSSION

Here, we reveal the increased susceptibility of *rve4 6 8* to osmotic and salt stress, along with the potential protein-level molecular mechanisms underpinning these responses, establishing a role for RVE8-like proteins in regulating plant drought-like stress responses. Through a combination of phenotypic, proteomic, and physiological analyses, we show that loss of RVE4, 6 and 8 resulted in reduced germination and decreased primary root elongation under osmotic and salt stress conditions, time-of-day dependent proteome remodeling and mis-regulation of photosynthetic and metabolic pathways. Finally, we provide analyses validating candidates related to fatty acid biosynthesis, vesicle trafficking and a receptor-like kinase related to ABA-related processes, as novel RVE- regulated proteins that impact the plant drought-like stress response.

### Proteomic analysis uncovers altered osmotic stress response in rve4 6 8 mutants

Given the increased osmotic and salt stress susceptibility exhibited by *rve4 6 8* plants, we hypothesized that we may observe altered proteome-level responses that contribute to this phenotype. Here, our GO (**Figure 3**) and association network (**Figure 4**) analyses identified proteome changes related to water deprivation response in *rve4 6 8* under both osmotic and salt stress conditions, particularly at ZT12 (end-of-day). To our surprise, many of these water deprivation response proteins exhibited higher abundance in *rve4 6 8*, particularly under osmotic stress (**Supplemental Figure 2**). This included COLD-REGULATED15A (COR15A; AT2G42450), EARLY RESPONSIVE TO DEHYDRATION 10 (ERD10; AT1G20450), LATE EMBRYOGENESIS ABUNDANT 7 (LEA7; AT1G52690), LEA14 (AT1G01470), LEA30 (AT3G17520), RESPONSIVE TO DESSICATION 28B (RD29B; AT5G52300), KIN1 (AT5G15960), RD20 (AT2G33380), RD22 (AT5G25610) and RD29A

(AT5G52310), and RD29B (AT5G52300) each demonstrating higher abundance at ZT12. By contrast under salt stress COR15A, RD20, RD22, RD29A and RD29B were of greater abundance in Col-0 at ZT12, emphasizing the differential impacts of osmotic versus salt stress in Arabidopsis (Rodriguez Gallo et al., 2023; Gonzalez et al., 2024), however, not in relation to the circadian clock from a proteome standpoint. A number of these genes have been implicated as having early transcriptional responses to osmotic stress (e.g., < 10 h following the osmotic stress onset (Kiyosue et al., 1994; Xiong et al., 2002; Narusaka et al., 2003), with our study revealing that proteome-level changes also manifest, rendering the results of this study of interest. Our proteomic analysis was performed after 8 days of osmotic stress; thus, the daily timing of water deprivation responses may be mis-regulated in *rve4 6 8*, conferring broader susceptibility to osmotic stress, as the plant is mounting too great of a stress response over a prolonged period of time. It would be interesting to assess the proteomic differences of *rve4 6 8* relative to Col-0 under osmotic and salt stress over time ranging from early responsive (e.g., hours) to days to determine whether the proteomic differences manifest as mistimed stress responses on a daily or multi-day timescale. However, given that the Col-0 osmotic stress related DAPs at ZT12 versus ZT0 were not significant in *rve4 6 8*, it is possible that the significant genotypic related DAPs we see in *rve4 6 8* versus Col-0 under osmotic stress is related to mis-regulated timing.

### The rve4 6 8 triple knockout shows inappropriate regulation of photosynthesis and photodamage responses under salt stress

Our proteomic analysis of *rve4 6 8* under drought-like stress conditions identified an altered abundance of proteins related to carbohydrate metabolism and photosynthesis, particularly under our salt stress conditions. We previously reported the starch excess phenotype of *rve4 6 8* at ZT0, rendering our results here an interesting finding given that it is more pronounced under abiotic stress (Scandola et al., 2022). To this point, association network analysis uncovered a strikingly higher abundance of photosynthesis and chlorophyll-related proteins in *rve4 6 8* under salt stress, particularly at ZT12 (**Supplemental Figure 2**), which we also observe manifesting as DAPs in Col-0 at ZT12 versus ZT0.

Here, we see a greater abundance of PHOTOSYSTEM I (PSA) subunits, PSAC (ATCG01060) and PSAO (AT1G08380), and PHOTOSYSTEM II (PSB) subunits, PSB28 (AT4G28660), PSBE (ATCG00580), PSBH (ATCG00710), PSBQ (AT4G21280), as well as LIGHT HARVESTING PROTEINS OF PHOTOSYSTEM I (LHCA) proteins, LHCA3 (AT1G61520), and minor antennae of photosystem II (LHCB), LHCB4.1 (AT5G01530), LHCB4.2 (AT3G08940), LHCB5 (AT4G10340), LHCB6 (AT1G15820), and LHCB7 (AT1G76570) in *rve4 6 8*. Additionally, we see a higher abundance of photosynthetic electron transport chain proteins including ATP SYNTHASE D CHAIN (ATPD; AT3G52300), NAD(P)H DEHYDROGRNASE (NDH) subunits, NDHI (ATCG01090), NDHS (AT4G23890), PHOTOSYNTHETIC ELECTRON TRANSFER C (PETC; AT4G03280), PETD (ATCG00730) as well as REDUCED INDUCTION OF NON-PHOTOCHEMICAL QUENCHING1 (RIQ1; AT5G08050) and SUPPRESSOR OF QUENCHING1 (SOQ1; AT1G56500), which are involved in non-photochemical quenching (Duan et al., 2023). Their expression at end-of-day (ZT12) and enrichment among the Col-0 DAPs at ZT12 versus ZT0, but not in *rve4 6 8*, suggest dysregulation of photosynthetic processes and ability to deal with photodamage responses. Plants typically downregulate photosynthesis under salinity stress (Zahra et al., 2022; Wijeweera et al., 2025), thus the increased susceptibility of *rve4 6 8* under these conditions may be due to mistimed photosynthetic responses versus an investment in stress mitigation effects. Our MultiSpeQ physiological measurements reinforce this hypothesis, as it found higher PhiNO in *rve4 6 8* relative to Col-0 under salt stress, suggesting greater photodamage through non-photochemical processes. Overall, our findings indicate an important role for RVE4, 6 and 8 in controlling the regulation of photosynthesis and photodamage responses under salt stress.

### Quantitative proteomics uncovers inappropriate timing of phenylpropanoid metabolism and anthocyanin accumulation in rve4 6 8 under salt stress

Phenylpropanoid metabolism is commonly reported as responsive to plant abiotic stress (Ninkuu et al., 2025). Our GO enrichment of phenylpropanoid metabolism amongst the Col-0 DAPs at ZT12 versus ZT0 under osmotic and salt stress, but not in *rve4 6 8* indicates dysregulation of phenylpropanoid metabolism in *rve4 6 8*. We additionally observed increased purpling in *rve4 6 8* plants, which we confirmed as greater anthocyanin accumulation under salt stress, suggesting altered regulation of anthocyanins and phenylpropanoid metabolism. Under salt stress, we see dysregulation of PHENYLALANINE AMMONIA-LYASE1 (PAL1; AT2G37040) and PAL2 (AT3G53260). We also observe FLAVONOL SYNTHASE5 (FLS5; AT5G63600) and CHALCONE SYNTHASE (CHS; AT5G13930) to have differential abundance in *rve4 6 8* versus Col-0. RVEs have been linked to appropriate timing of anthocyanin accumulation in pear fruit peel (Li et al., 2020; Li et al., 2024) and RVE8 binds promoters of anthocyanin biosynthesis genes in Arabidopsis to control the correct daily timing of accumulation (Pérez-García et al., 2015). Thus, it is likely that *rve4 6 8* is inappropriately timing the production of anthocyanin metabolism proteins, signaling increased stress and supporting our observations of anthocyanin accumulation under salt stress.

### Phenotypic validation of putative RVE8-like protein-controlled targets involved in abiotic stress response

We selected three proteins related to biological processes altered in *rve4 6 8* plants in response to abiotic stress to validate our dataset. All three proteins are related to abiotic stress responsive pathways but had not been fully characterized as associated with either osmotic or salt stress response. This included: LACS3, SYP43 and CST, which are involved in fatty acid biosynthesis, vesicle transport and ABA-related processes, respectively. Our validation experimentation included assessing *lacs3, syp43* and *cst* plants for altered root length under osmotic and salt stress (**Figure 6**). Each gene-deficient plant line possessed increased root length under salt stress, suggesting they may each be abiotic stress susceptibility factors. Interestingly, we did not see a phenotype under the applied mannitol stress; however, we used the lower mannitol concentration where we observed a phenotype in *rve4 6 8*, and thus an increased concentration may elicit a stronger phenotype given the wide-ranging control RVE8-like proteins have over a diversity of drought-related proteins and processes, as evidenced by our proteome data.

We first examined LACS3, a protein responsible for very-long-chain fatty acid biosynthesis (Pulsifer et al., 2012). Many LACS genes are related to jasmonic acid signaling (Kienow et al., 2008), which we observed enriched among proteins of higher abundance in Col-0 under salt stress (**Figure 4B**). Thus, LACS3 may have a link to jasmonic acid signaling and its control through RVE8-like proteins. We additionally observed a higher abundance of vesicle transport proteins, which are known to be important for salt stress responses (Garcia de la Garma et al., 2015), under both osmotic and salt stress in *rve4 6 8*. Therefore, we selected one of the highest DAPs, SYP43, for further analysis. Several SYPs have been implicated in abiotic stress responses (Baena et al. 2024; Chen et al., 2019), with SYP43 having not been previously identified as abiotic stress responsive. However here, *syp43* exhibited higher tolerance to salt stress, maintaining a longer primary root length (**Figure 6**), suggesting that *rve4 6 8* may have mis-regulated vesicle transport. Finally, one of the highest changing DAPs in *rve4 6 8* relative to Col-0 under both osmotic and salt stress conditions was CST, an RLK with a key role in abscission (Burr et al., 2011; Groner et al., 2016). Given the relationship to abiotic stress, this connects RVE8-like proteins to ABA-related responses, of which abscission is closely linked (Patharkar and Walker, 2016). ABA has been connected to reduced leaf abscission and salt tolerance in citrus plants (Gómez-Cadenas et al., 2002), rendering our finding of salt stress in agreement. Given that CST is an RLK, future research should determine putative phosphorylation targets for CST under salt stress.

### Conclusions

Here, through a combination of phenotyping and quantitative proteomics, we have revealed the molecular underpinnings for how RVE8-like proteins in Arabidopsis impact osmotic and salt stress. The *rve4 6 8* plants possess extensive perturbations in osmotic stress responsive proteins under mannitol treatment and increased abundance of photosynthesis proteins, particularly at end-of-day, under salt stress conditions, suggesting altered regulation of photosynthesis. Additionally, we find phenylpropanoid metabolism perturbed in *rve4 6 8*, suggestive of a heightened overall stress response state that leads to susceptibility under drought-like stress conditions. Finally, using a series of gene-deficient genetics for some of the highest changing DAPs related to the loss of RVE8-like proteins, we validate three putative RVE8-like protein-regulated targets involved in fatty acid metabolism, vesicle transport and abscission.

With our study limited to total proteome abundance responses, but revealing large changes in proteins such as CST, future global analysis of post-translational protein modification (PTM) changes, such as phosphorylation, would further enrich our understanding of the *rve4 6 8* stress-responsive proteome changes. Overall, our results indicate an important role for RVE8-like proteins in regulating the plant responses to osmotic and salt stress, providing further link for the circadian clock to important agricultural abiotic stresses responses.

## MATERIALS AND METHODS

### Plant growth and validation under osmotic and salt stress conditions

For phenotyping experiments, Col-0 (wild-type) and *rve4 6 8* seeds were rinsed in 70% (v/v) ethanol for 2 min, followed by a 30% (v/v) bleach (Clorox® 7.5%) wash for 7 min, and final three consecutive washes with sterile water. The seeds were then imbibed on 0.5x MS media, pH 5.8 with 7 g/L agar (Sigma). All seeds were stratified for 3 days at 4 °C in the dark and then exposed to 12 h light: 12 h dark regime. For root length experiments, seeds were germinated and grown on 0.5x MS plates for 5 days before transfer to experimental plates. Experimental conditions consisted of 0.5x MS plates as control, with 50 mM, 100 mM or 150 mM mannitol, or with 25 mM, 50 mM or 100 mM NaCl for 8 days. For germination experiments, seeds were sown directly on experimental plates: Control (0.5x MS), 300 mM Mannitol or 100 mM NaCl and germination was measured over 10 days at 22 °C. On the fourth day or the last day of the time course for germination and root length experiments, respectively, photographs of the seedlings were taken, and primary root length measurements were taken.

### MultispeQ and anthocyanin measurements on salt-treated plants

Col-0 and *rve4 6 8* plants seedlings were grown for 21 days on soil and watered as normal. After 21 days, plants were watered every other day with 300 mM NaCl for one week for salt stress treatment.

Photosynthetic parameters were measured using the MultispeQ v1.0 device (PhotosynQ LLC, USA; (Kuhlgert et al., 2016) following the Photosynthesis RIDES 2.0 protocol. Plant leaves were clamped at ZT3 ensuring full coverage. The leaf was clamped for ∼10 s while the collection process was completed. This was done for a total of n=16 plants per condition. At the same stage, leaves were collected for anthocyanin analysis, performed as previously described (Neff and Chory, 1998). Briefly, 500 µL of 100% (v/v) methanol- 1% (v/v) hydrochloric acid was added to 100 mg of ground leaf material and incubated in darkness for 24 h at 4 ℒ. Next, 200 µL of sterile water and 500 µL of chloroform was added followed by room temperature centrifugation at 18, 000 x g for 5 min. Supernatant (400 µL) was transferred to new tubes and 400 µL of 60% (v/v) methanol-1% (v/v) HCl was added. From this, 200 µL from each sample was transferred into a 96 well plate and absorbance was measured at 530 nm and 657 nm with a Tecan Spark® plate reader using the 60% (v/v) methanol – 1% (v/v) HCl as blank. Relative anthocyanin content was calculated as follows: Relative anthocyanin content= (Abs_530_- Abs_657_) x 1000 x powder weight (mg)^-1^.

### Plant growth and tissue harvesting for quantitative proteomics

Col-0 and *rve4 6 8* seeds were sterilized, stratified, germinated, and transferred as described above after 5 days and grown on experimental plates (Control, 50 mM mannitol, or 100 mM NaCl) for 8 days. Whole seedlings were harvested at zeitgeber time 0 (ZT0) and ZT12 and immediately flash-frozen in liquid nitrogen. Samples were stored at -80 °C prior to their use for quantitative proteomics and downstream data analyses.

### Tissue processing, protein extraction and digestion

Tissue was extracted at a 1:2 (w/v) ratio with a solution of 50 mM HEPES-KOH pH 8.0, 50 mM NaCl, and 4% (w/v) SDS. Samples were vortexed and placed at 95 °C in a table-top shaking incubator (Eppendorf) at 1,100 x g for 15 min, followed by an additional 15 min shaking at room temperature. Samples were then clarified at 20,000 x g for 5 min and supernatant was transferred to fresh 1.5 mL Eppendorf tubes. Sample protein concentration was determined by bicinchoninic acid (BCA) assay (23225; Thermo Scientific). Sample reduction was performed using 10 mM dithiothreitol (DTT) at 95 °C for 5 min, followed by cooling to room temperature and alkylation with 30 mM iodoacetamide (IA) for 30 min in darkness at room temperature without shaking. Subsequently, 10 mM DTT was added to each sample, followed by a quick vortex, and then incubation at room temperature for 10 min without shaking. Total proteome peptide pools were then generated using a KingFisher Apex (5400910; Thermo Scientific) automated sample preparation device as outlined by Leutert et al. (2019) without deviation. Sample digestion was performed using sequencing grade trypsin (V5113; Promega), with generated peptide pools quantified by Nanodrop and acidified with formic acid (FA) to a final concentration of 5% (v/v) prior to being desalted using 1cc tC18 Sep Pak cartridges (WAT036820; Waters) as previously described (Scandola et al., 2022; Uhrig et al., 2019). All peptides were then dried by vacuum centrifugation and resuspended in 3% (v/v) ACN / 0.1% (v/v) FA immediately prior to MS analysis.

### LC-MS/MS analysis

Protein abundance changes were assessed using a FAIMS mounted Fusion Lumos Tribrid Orbitrap mass spectrometer (Thermo Scientific) in data independent acquisition (DIA) mode using the BoxCarDIA method (Mehta et al., 2022; Scandola et al., 2022; Rodriguez Gallo et al., 2023). Dissolved peptides (1 µg) were injected using an Easy-nLC 1200 system (LC140; Thermo Scientific) and separated on a 50 cm Easy-Spray PepMap C18 column (ES803A; Thermo Scientific). A spray voltage of 2.2 kV, funnel RF level of 40 and heated capillary at 300 °C was deployed, with all data acquired in profile mode using positive polarity, with peptide match turned off and isotope exclusion selected. All gradients were run at 300 nL/min with the analytical column temperature set to 50 °C. Peptides were eluted using a segmented solvent B gradient of 0.1% (v/v) FA in 80% (v/v) ACN from 4-41% B (0-107 min). FAIMS using compensation voltages (CVs) of -30, -50, -70 were used with static gas flow rate of 3.5 L/min. Within each CV, BoxCarDIA acquisition was performed as previously described (Mehta et al., 2022; Scandola et al., 2022; Rodriguez Gallo et al., 2023). MS1 analysis was performed by using two multiplexed targeted SIM scans of 10 BoxCar windows each, with detection performed at a resolution of 120,000 at 200 m/z and normalized AGC targets of 100% per BoxCar isolation window. Windows were custom designed as previously described (Mehta et al., 2022; Scandola et al., 2022; Rodriguez Gallo et al., 2023). An AGC target value for MS2 fragment spectra was set to 2000%. Twenty-eight 38.5 m/z windows were used with an overlap of 1 m/z. Resolution was set to 30,000 using a dynamic maximum injection time and a minimum number of desired points across each peak set to 6.

### LC-MS/MS data analysis

All acquired BoxCarDIA data was analyzed in a library-free DIA approach using Spectronaut v16 (Biognosys AG) using default settings. Key search parameters included: a protein, peptide and PSM FDR of 1%, trypsin digestion with 1 missed cleavage, fixed modification including carbamidomethylation of cysteine residues and variable modification including methionine oxidation. Data was Log2 transformed, globally normalized by median subtraction with significantly changing differentially abundance proteins determined and corrected for multiple comparisons (Bonferroni-corrected p-value ≤ 0.05; q-value ≤ 0.05).

### Bioinformatics and data visualization

Gene Ontology (GO) analyses were undertaken using The Ontologizer (http://ontologizer.de/) using a parent-child enrichment analysis (p-value < 0.01). Visualization was done using R and R package ggplot2 in combination with Affinity Designer. All data were plotted using GraphPad Prism (version 8).

Association networks were created in Cytoscape version 3.10.4 using the STRING-DB and enhancedGraphics Cytoscape application using a minimum edge threshold of <0.7. Metabolic pathway enrichment analysis was undertaken using the Plant Metabolic Network (PMN; https://plantcyc.org/) using a test threshold of p-value < 0.01 to determine enrichment.

## ACKNOWLEDGEMENTS

The authors thank Dr. Stacey Harmer for providing the *rve4 6 8* and corresponding Col-0 seeds. We would also like to thank G2V Optics Inc. for providing their LED lighting system. We additionally would like to thank Jack Moore of the Alberta Mass Spectrometry and Proteomics Facility. The authors thank the Natural Sciences and Engineering Research Council (NSERC) and Canada Foundation for Innovation (CFI) for funding.

## AUTHOR CONTRIBUTIONS

LEG: Investigation; Formal Analysis; Methodology; Data Curation and Visualization; Writing and Editing.

IK: Investigation; Methodology SS: Investigation; Methodology DM: Investigation; Methodology

RGU: Conceptualization; Methodology; Supervision; Project Administration; Data Curation; Writing and Editing; Funding Acquisition

## SUPPLEMENTARY DATA

**Supplementary Figure S1.**
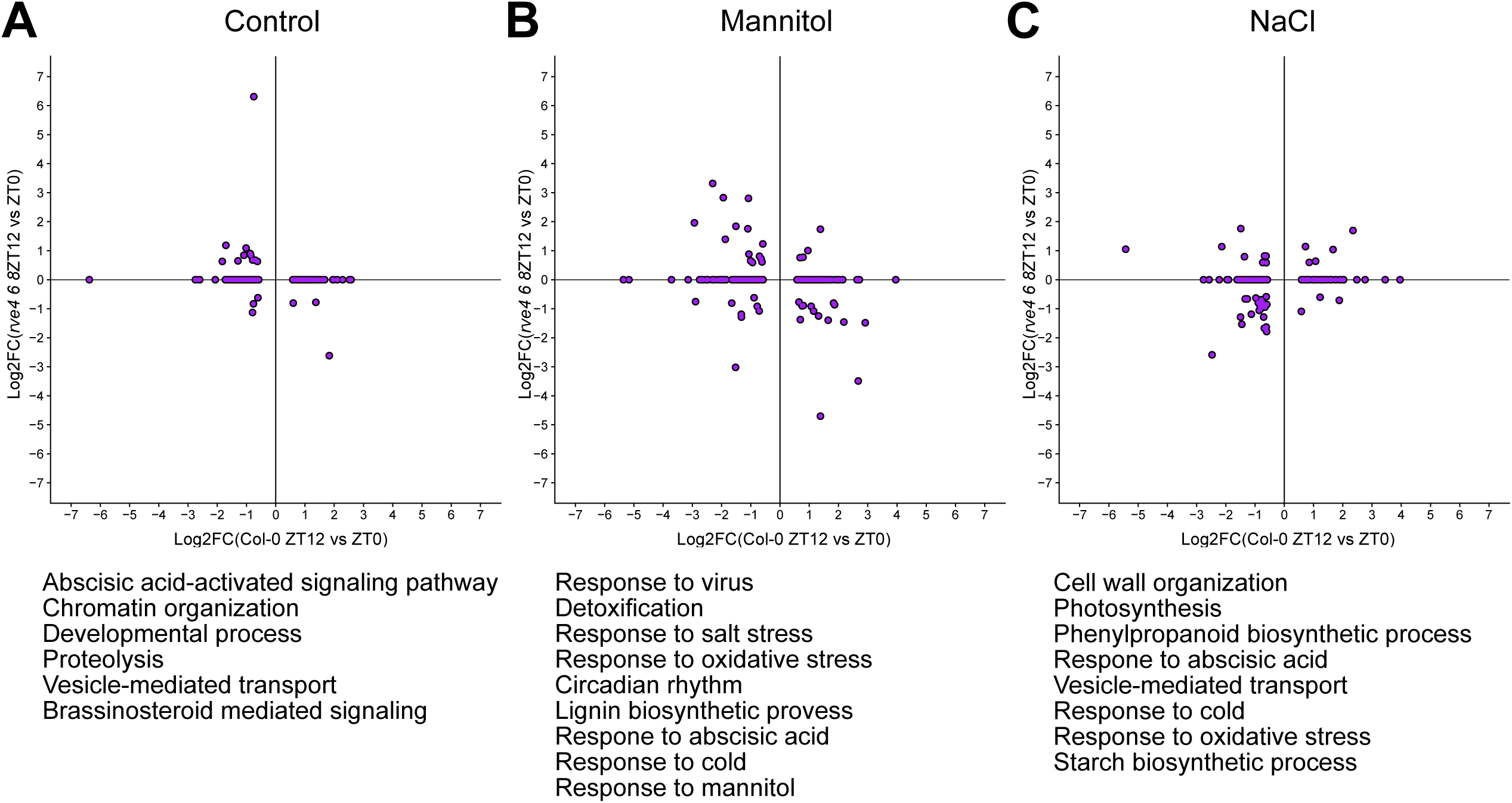
Distribution of significantly changing proteins (ZT12 vs. ZT0) under Control (**A**), Mannitol (**B**) and NaCl (**C**) treatments (Log2FC > 0.58 or < -0.58). Below each graph are enriched GO terms for those proteins that are significantly different in Col-0 between the two sampling timepoints, but not enriched in *rve4 6 8* under the same condition.

**Supplementary Figure S2.**
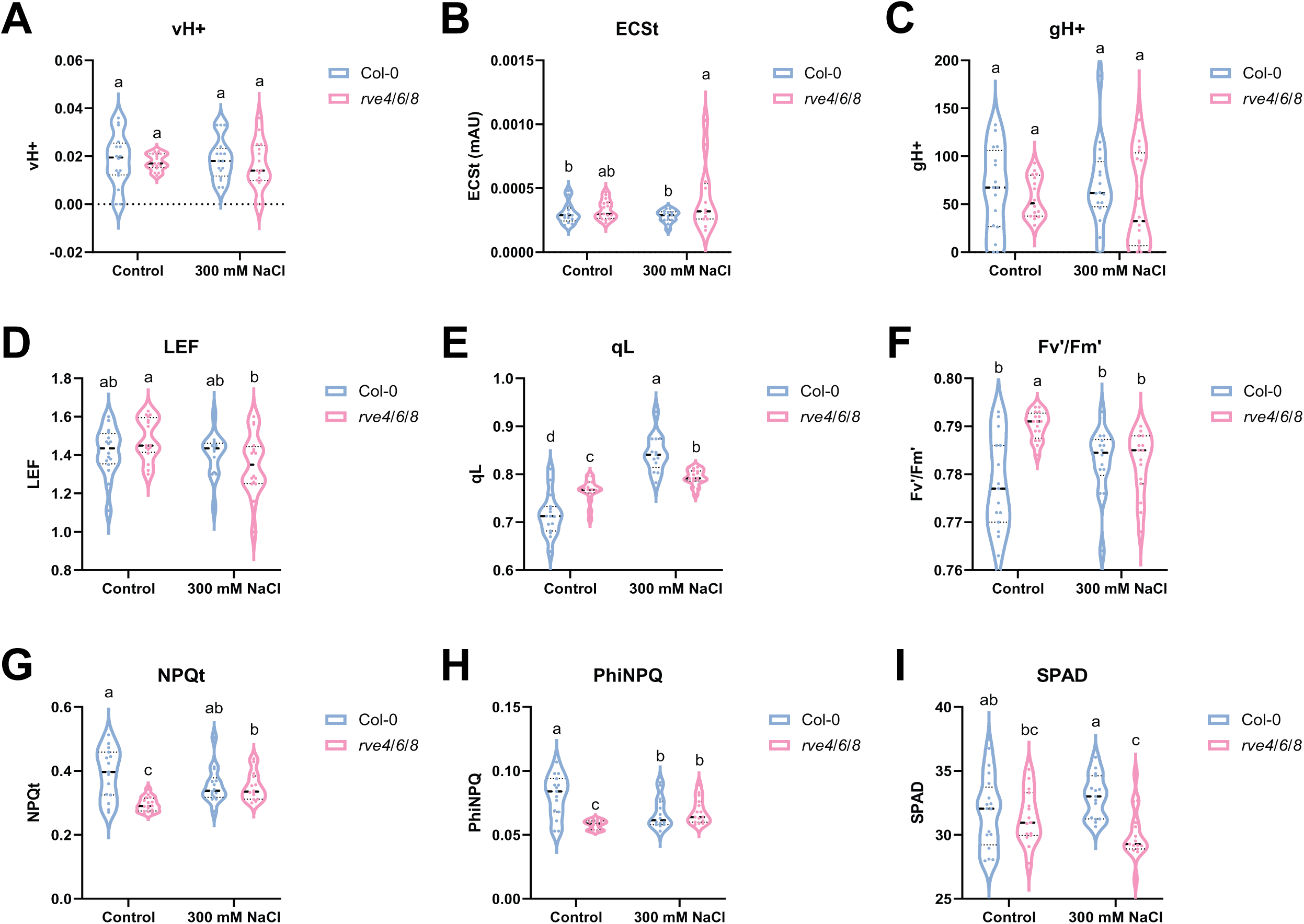
MultispeQ parameters of *rve4 6 8* mutant compared to Col-0 under control and NaCl conditions. Violin plots indicating photosynthetic parameters of the plants grown under control and salt conditions as described in Figure 5 for Col-0 (blue) and *rve4 6 8* mutant (pink) under control and 300 mM NaCl. **A)** steady-state proton flux across the thylakoid membrane (vH+), **B)** Electrochromic Shift (ECSt), **C)** proton conductivity of thylakoid membrane (gH+), **D)** linear electron flow (LEF), **E)** photochemical quenching (qL), **F)** Efficiency of photosystem II (Fv’/Fm’), **G)** Non-photochemical quenching (NPQt), **H)** Quantum yield of non-photochemical quenching (PhiNPQ), **I)** Relative chlorophyll (SPAD). Violin plots indicate the median and n = 16 for photosynthesis measurements and 4 for anthocyanin. Letters denote statistical significance (p < 0.05) based on a 2-way ANOVA followed by Fisher’s LSD test for multiple comparisons.

**Supplementary Figure S3.**
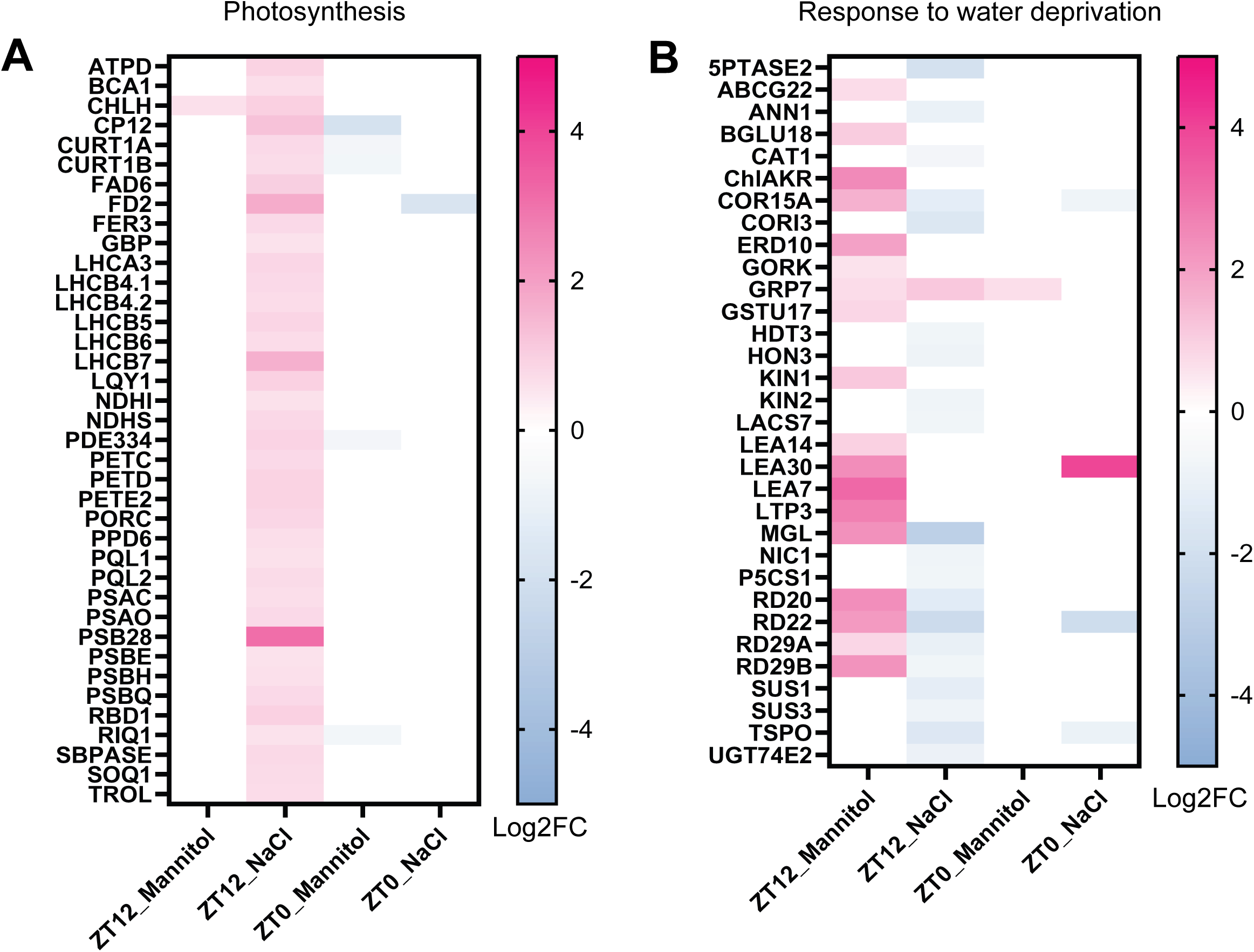
Heatmap summary of significantly changing proteins in *rve4 6 8* that are involved in Photosynthesis (GO:0015979) (**A**) and Response to Water Deprivation (GO:0009414) (**B**). Values from blue (less abundant) to pink (more abundant) depict the Log2FC change in *rve4 6 8* relative to Col-0.

**Supplementary Table S1. All quantified and significant proteins.**

**Supplementary Table S2. AraCyc Plant Metabolic network enrichment.**

